# Real-time individual benefit from social interactions before and during the lockdown: The crucial role of personality, neurobiology and genes

**DOI:** 10.1101/2021.06.02.446719

**Authors:** Maximilian Monninger, Pascal-M. Aggensteiner, Tania M. Pollok, Iris Reinhard, Alisha S.M. Hall, Lea Zillich, Fabian Streit, Stephanie-H. Witt, Markus Reichert, Ulrich Ebner-Priemer, Andreas Meyer-Lindenberg, Heike Tost, Daniel Brandeis, Tobias Banaschewski, Nathalie E. Holz

**Affiliations:** Department of Child and Adolescent Psychiatry and Psychotherapy, Central Institute of Mental Health, Medical Faculty Mannheim / Heidelberg University, J5, Mannheim 68159, Germany; Department of Biostatistics, Central Institute of Mental Health, Medical Faculty Mannheim / Heidelberg University, J5, Mannheim 68159, Germany; Department of Genetic Epidemiology in Psychiatry, Central Institute of Mental Health, Medical Faculty Mannheim / Heidelberg University, J5, Mannheim 68159, Germany; Department of Psychiatry and Psychotherapy, Central Institute of Mental Health, Medical Faculty Mannheim / Heidelberg University, J5, Mannheim 68159, Germany; mental mHealth lab, Institute of Sport and Sports Science, Karlsruhe Institute of Technology, Engler-Bunte Ring 15, 76131 Karlsruhe, Germany; Department of Child and Adolescent Psychiatry and Psychotherapy, Psychiatric Hospital, University of Zurich, Neumünsterallee 9, Zurich 8032, Switzerland; Center for Integrative Human Physiology, University of Zurich, Winterthurerstr. 190, Zurich 8057, Switzerland; Neuroscience Center Zurich, University of Zurich and ETH Zurich, Winterthurerstr. 190, Zurich 8057, Switzerland

**Keywords:** Ecological momentary assessment, longitudinal study, neuroimaging, social interactions, COVID-19

## Abstract

**Background:** Social integration is a major resilience factor for staying healthy. However, the COVID-19-pandemic led to unprecedented restrictions in social life. The consequences of these social lockdowns on momentary well-being are yet not fully understood.

**Method:** We investigated the individual affective benefit from social interactions in a longitudinal birth cohort study. We used two real-time, real-life ecological momentary assessments once before and once during the initial lockdown of the pandemic (N~6800 total observations) to determine the protective role of social interactions on well-being. Moreover, we used a multimethod approach combining the ecological assessment data with individual risk and resilience factors to analyze the moderating mechanisms of personality, neurobiology and genes.

**Results:** Social contacts were linked to higher positive affect both during normal times and during the COVID-19 pandemic, highlighting the beneficial role of social embedding. Moreover, this relationship was moderated by amygdala volume, neuroticism and polygenic risk for schizophrenia. In detail, participants with a larger left amygdala volume and higher trait neuroticism exhibited an affective benefit from more social interactions prior to the pandemic. This pattern changed during the pandemic with participants with smaller amygdala volumes and lower neurotic traits showing a social affective gain during the pandemic. Moreover, participants with low genetic risk for schizophrenia showed an affective benefit from social interactions irrespective of the time point.

**Conclusion:** Our results highlight the protective role of social integration on momentary well-being. Thereby, we offer new insights into how this relationship is differently affected by a person’s, neurobiology, personality, and genes under adverse circumstances.

## Introduction

At the beginning of 2020, the coronavirus (COVID-19) outbreak was declared a global pandemic by the World Health Organization (WHO). In addition to posing a dramatic public health burden, the pandemic also brought drastic social contact restrictions (“*lockdown”)*, which began on the 23^rd^ of March 2020 in Germany. During this lockdown period, individuals were only allowed to meet with people from one other household. First studies from China investigating the mental health outcomes of the pandemic demonstrated elevated levels of stress, anxiety, and depressive symptoms (1, 2). However, the specific impact of social contact restrictions on mental health remains largely unknown. Therefore, the current study aims to investigate the relationship between social contacts and well-being before and during the lockdown phase in Germany using a real-time, real-life approach in the framework of a longitudinal study.

Social integration plays an important role in promoting resilience. The amount of social interactions, the social network size (i.e., the number of people with whom an individual is usually in contact with), and self-perceived social support are linked to subjective well-being (3–6), whereas a lack of social integration and the apprehended feeling of loneliness are associated with poorer mental (7) and physical health (8–10). A recent study using ecological momentary assessment (EMA) found that participants who reported recent social interactions experienced more positive mood (11). However, little is known about how social contact restrictions affect momentary affective states. To our knowledge, there is only one published study comparing social network characteristics prior to and during the lockdown phase from a longitudinal perspective: In a Swiss student sample, participants reported significantly fewer interactions and study partners on average during than before the lockdown, although friendships and perceived social support did not change (12), indicating that structural characteristics (i.e., the objective quantity of social interactions) are more impaired due to the lockdown than evaluative and subjective characteristics of social relationships (i.e., the self-perceived quality of a social interaction). Critically, this study did neither address the momentary effect of social interactions on affective states, nor explore possible moderators of this relationship.

However, so far there is a lack in understanding the underlying mechanisms of the beneficial relationship between social integration and affective well-being. Therefore, it is important to consider an individuals’ risk and resilience factors, including neurobiology, personality and genetic make-up, all of which determine how much an individual might profit from its social embedding. As an example, the amygdala is a core structure of the social brain (13). It has been shown to be affected by socioenvironmental influences (14) and plays a particularly prominent role in how a person is socially integrated. As such, previous studies found a positive relationship between larger amygdala volumes and increased social network sizes (15, 16), heightened perceived social support (17), and social connectedness (18). Further, in terms of personality, high levels of neuroticism are correlated with lower frequency of involvement in real-life social interactions, a smaller social network and an overall weaker social bonding (19). In addition, neuroticism was linked to self-reported changes in social behavior due to the COVID-19-pandemic (20). Moreover, the genetic risk for schizophrenia has been highlighted as conferring a maladaptation to social contexts (21, 22), with patients suffering from schizophrenia often characterized by weak social integration, a reduced social network size and fewer friends (23, 24).

The COVID-19 lockdown offers a unique opportunity to study risk and resilience factors in a naturalistic, real-life framework. Critically, the above mentioned neurobiobehavioral markers have all been related to how well individuals adapt to adverse events. For instance, the amygdala has proven as key convergence site of social adversity (25) and is critically involved in stress adaptation (25–28). In addition, neuroticism is linked to elevated levels stress reactivity, heightened risk of mood disorders, and overestimation of potential health threats (29–31). Moreover, higher neurotic traits significantly predicted decreased emotional, psychological and social well-being during the COVID-19 pandemic (20). Similarly, schizophrenia has been previously associated with higher affective reactivity to daily stressors (32), suggesting a heightened sensitivity to the COVID-19 pandemic in individuals with a higher polygenic risk for schizophrenia (SCZ-PRS).

Thus, due to their involvement in risk and resilience pathways, these neurobiobehavioral markers are specifically qualified to modulate how social interactions promote well-being before and during the pandemic. Within the framework of an ongoing longitudinal birth-cohort at risk (“Mannheim Study of Children at Risk”), we assessed neurobiobehavioral moderators prior to the pandemic and social interactions along with well-being using EMA before and during the lockdown. Based on previous findings (11) and on the protective role of social interactions when adversity is encountered (14), we expected to find a strong positive association between social interactions and well-being both prior to and during the pandemic. Moreover, our approach allowed us to further investigate the moderating effects of amygdala volume, neuroticism, and SCZ-PRS on the impact of social interactions on momentary well-being. Given the role of the amygdala, neuroticism, and SCZ-PRS in socioenvironmental risk and resilience (15, 21, 29, 32–34), we expected individuals with larger amygdalae volumes, lower neurotic traits and lower SCZ-PRS to show an affective benefit from social interactions and tested whether this is different under social contact restrictions.

## Materials and Methods

### Sample

The present investigation was conducted in the framework of the Mannheim Study of Children at Risk (‘MARS’), an ongoing prospective study of the long-term outcomes of early psychosocial and biological risk factors following children since birth. The initial sample consisted of 384 children born between 1986 and 1988 in the Rhine-Neckar region of Germany and were included according to a two-factorial design intended to enrich and control the risk status of the sample (see (35) for full details).

Starting at the age of 3 months, information on mental health, personality traits, and genetic variability was collected prospectively up to the most recent assessment wave at the age of 32-33 years, which was disrupted by the COVID-19 pandemic. This assessment consisted of a questionnaire package on physical and mental health, a diagnostic interview, MRI measurements, and an EMA. Starting shortly after the social contact restrictions were put in place in Germany in April 2020, participants who had completed the EMA week (n=165) were invited to take part in an online survey and to repeat the EMA procedures during the COVID-19 pandemic. A total of 133 participants completed the online survey and 70 participated (distribution in the current sample: 22 (31.4 %) participants without psychosocial risk, 26 (37.1%) with low psychosocial risk, and 22 (31.4%) with high psychosocial risk at birth) in both EMA measurements. The study was approved by the Ethics Committee of the University Heidelberg, Germany, written informed consent from all participants was obtained, and participants were financially compensated.

### EMA procedures

Participants were asked to install a commercial e-diary app (MovisensXS, version 1.4.3) on their own Android smartphone. The e-diary started on a fixed date, with the participants receiving prompts via an acoustic, visual, and vibration signal. Prompts were scheduled from 8am until 10pm with a fixed interval of 120minutes to facilitate retrospection, resulting in a maximum of eight prompts per day and 56 prompts per week. Upon receiving a prompt, participants completed the questionnaire, which took approximately 90seconds. Participants had the opportunity to postpone a prompt for a maximum of 25minutes. The same procedure was repeated during the lockdown phase of the COVID-19 pandemic in Germany beginning on the 23^rd^ April 2020 (Supplementary Figure 1).

### Affective state

Positive and negative affect was measured using a 15-item short version of the German adaptation of the Positive and Negative Affect Schedule (PANAS) (36) with additional items capturing stress reactivity (37–40). Participants were asked to rate their current positive or negative feelings on a 7-point Likert scale (1=fully disagree, 7=fully agree). Mean scores for positive and negative affect were calculated for each prompt and used in all analyses as dependent variables. Between- and within-person reliability coefficients for positive (R_kf_=0.99; R_cn_=0.62) and negative affect (R_kf_=0.99; R_cn_=0.75) were calculated using mixed models(41) and ranged from moderate to high.

### Momentary social contacts

Participants were asked to indicate the number of real-life social contacts and the quality of the most important interaction within the last two hours before the prompt (i.e., the interval between two prompts). Real-life social contacts are defined as interactions in which participants were talking to or interacting with another person face-to-face. For the quality rating, participants were asked to indicate on a visual analogue scale ranging from 0 to 100 how positive the most important interaction was experienced (0 = very negative, 100 = very positive). If participants reported no interactions within the current time-frame, no follow-up question was presented.

### Stress burden during COVID-19

We used two items rated on a 10-point Likert scale (0: very low; 10: very high) to assess the impact of COVID-19 on physical and mental health (*‘The physical burden of COVID-19 for me is…’, ‘The mental burden of COVID-19 for me is…’*).

### Moderator variables

We tested for a possible moderating impact of the bilateral amygdala volume (n=70), neuroticism (n=69), and SCZ-PRS (n=68) on the relationship of social interactions with positive and negative affect.

### Amygdala volume

At the age of 32–33 years and prior to the COVID-19 pandemic, high-resolution anatomical images with 208 slices covering the whole brain were acquired using a 3T-scanner (PrismaFit; Siemens) with a standard 32-channel head coil. Volumetric segmentation was performed with the FreeSurfer image analysis suite (Version 6.0.0) as described previously (42) to indicate left and right amygdala volume (mm^3^).

### Personality traits

Neuroticism was assessed at the age of 25 years using the German version of the NEO Five-Factor Inventory (NEO-FFI) (43), a widely used instrument to determine the Big Five personality traits. It contains a total of 60 items, with 12 items for each personality trait, rated on a 5-point Likert scale (1=strongly disagree, 5=strongly agree). Sum scores for neuroticism were calculated, with higher values representing a higher trait expression.

### Polygenic risk scores

DNA was extracted from whole blood or saliva of 306 participants of the initial sample. Genome-wide genotyping was performed using Global Screening Array 24 version 2 (Illumina, Inc., San Diego, CA, USA) at the Life & Brain facilities, Bonn, Germany. Quality control and filtering was performed using PLINK v1.90b6.7 (44), removing participants with >.02 missingness, heterozygosity rate >|.20|, and sex-mismatch. SNPs with a minor allele frequency of <.01, deviating from Hardy-Weinberg equilibrium (HWE) with a p-value of <10^−6^ and missing data >.02 were removed. Relatedness and population structure were filtered based on a SNP set filtered for high quality (HWE P>.02, MAF>.20, missingness = 0), and LD pruning (r^2^=.1). If subjects were cryptically related (pi hat>.10), one subject was excluded at random. Control for population stratification was performed by generating principal components and outliers, defined as deviating more than 4 SD on one of the first 20 principal components were excluded. Quality control and filtering resulted in a data set of 301 individuals and 482,981 SNPs. Of those, 68 subjects with available EMA data were included in the present analyses.

SCZ-PRS were calculated based on the genome-wide association data of 77,096 individuals (33,640 cases, 43,456 controls) from the Psychiatric Genomics Consortium (PGC) for SCZ (45). SCZ-PRS were calculated for 68 participants of the present study with PRSice version 2.2.6 (46) after clumping SNPS (linkage disequilibrium r^2^ < 0.1 250kb sliding window) for multiple p-value thresholds. For the analysis in the present study, we selected the PT=0.05, as this was the threshold with the best prediction in the discovery samples.

### Covariates

Gender, time of day, estimated intracranial volume (ICV), and the first ten principal components (to control for population stratification) were included as covariates when applicable. To ensure that the level of stress-dependent changes in positive affect were not due to biological programming by early postnatal stressful environments or encountered stress during lifetime, we additionally controlled for psychosocial risks at birth and stressful life events over the lifespan.

### Psychosocial risk

Psychosocial risk was assessed using a standardized interview according to an enriched family adversity index (47) at the participants’ age of 3 months, covering 11 items of the family environment, the parents, and their partnership (e.g., parental psychiatric disorders, overcrowding, or ongoing parental conflicts). A sum score of psychosocial risks were calculated by adding up the presence of all items.

### Stressful life events

Life events were recorded using a modified version of the Munich Events List (MEL;(48)) starting at the first assessment wave at the age of 3 months until the last assessment wave prior the COVID-19 pandemic. The MEL covered several areas of acute and chronic, positive and negative stressors, which were adjusted for different developmental stages. (e.g., school entrance at the age of 8 years; university entrance at the age of 19 years, but also chronic illness of a relative, ongoing parental disharmony or loss of a family member). Sum scores for each assessment wave were added to calculate an overall life events score.

### Data analysis

A-priori performed power calculations revealed 80% power to detect medium-sized effects with a compliance rate above 85 % based on a sample size of 70 participants with 14 observation days and a maximum of 8 responses per day.

Multilevel analyses were conducted to analyze the association between the quantity of social interactions (i.e. the number of social interactions) and current affective states as well as the statistical interaction with time point (pre- and during COVID-19 pandemic) and the potential moderators amygdala volume, neuroticism, and SCZ-PRS. Momentary affective states acted as the dependent variable and the number of social contacts as predictor variable (level-1), which were person-mean centered and nested within participants (level-2). Amygdala volume and neuroticism as level-2 variables were grand-mean centered. In addition, for visualization purposes, SCZ-PRS were z-standardized. To investigate the impact of COVID-19, a dichotomous time point variable (0 = pre-COVID-19, 1 = during-COVID-19) was included in all multilevel models. Furthermore, covariates of no interest, consisting of gender, time of day, psychosocial risk factors, ICV, and the first ten principal components of population stratification, when applicable, were entered in all models. Psychosocial risk factors, stressful life events, and ICV were grand-mean centered, whereas time of day was calculated in hours by subtracting the daily start time (i.e., 8am) from all values. In addition, the aggregated person-means for real-life social contacts were entered in all models separately to control for their potential effects.

We fitted five mixed models including random intercepts as well as random slopes for all level-1 predictors. In Model-I, we included real-life social contacts and time point as main effects of interest, and the corresponding two-way interaction effects. Models-II to V were based on Model-I, but additionally included one of the potential moderators at a time, i.e. left or right amygdala volume, neuroticism, or SCZ-PRS as a main effect, together with the corresponding two-way and three-way interaction effects. In a sub-analysis, we specified the effect of the quality of the most important interaction as an additional predictor (level-1, person-mean centered) for momentary affect with the corresponding two-way and three-way interaction effects. Given that the quality of social interaction was only assessed if social interactions have been indicated, resulting in fewer observations (N = 5820), this model had a lower power and is thus considered exploratory (see Supplement for further analyses with the quality of social interactions as additional predictor). All multilevel models were designed with the freely available R packages lme4 (49), and lmerTest (50) to compute p-values. To further analyze the interaction effects, simple slope analyses and Johnson-Neyman plots were computed in order to estimate the range of values of the moderator variable in which the slope of the predictor is significant vs. nonsignificant. For all analyses, the two-sided alpha level was Bonferroni-corrected set at 0.01 (p = 0.05 divided by 5 models).

## Results

### Descriptive data

In total, 6837 prompts (mean prompts per person 97.9, SD = 12.18; range = 57-112) were answered by 70 participants (63 % female; mean age = 33.36 years), resulting in a high compliance rate of 87.41% across both time points (baseline: mean = 49.86 prompts, COVID-19: mean = 48.04 prompts, Supplementary Table 1). The earliest start date of the assessment was 61.43 weeks prior to the social contact restrictions in Germany (23^rd^ March 2020) for the baseline data (latest inclusion date for baseline data 2^nd^ March 2020) and 4.57 weeks (32 days) after the restrictions were put in place for the COVID-19 assessment. As expected, participants reported significantly more weekly real-life contacts (mean=3.89; SD=2.81) in the baseline assessment compared to the COVID-19 assessment (mean=2.50; SD=1.37; t_138_=3.734, P<.001) while the quality of interactions (mean=76.15; SD=18.27, range=0-100) did not change (P=0.81). Given that only the quantity of social interactions changed between time points, we focused on this variable in the moderation analyses.

Overall, while positive affect decreased during COVID-19 (P<0.001), momentary negative affect was higher (P=0.005). Additional linear regression analyses revealed that COVID-19-related stress predicted lower positive affect (P=0.038). This did not pertain to social contact reductions (P=0.664) or negative affect (P > 0.8).

### Social contacts and affective state (Model-I)

The number of real-life contacts and the quality of the most important interaction were significantly associated with positive affect across both time points (P=0.001; Supplementary Table 2a&2b, Figure 1), indicating a mood-uplifting effect by social interactions. In addition, a more pleasant social interaction was associated to lower negative affect (P<0.001, Supplementary Table 3b), however, no such relationship was found regarding the quantity of real-life contacts (P=0.540; Supplementary Table 3a). Therefore, no further analyses with negative affect as outcome were performed.

**Figure 1:**
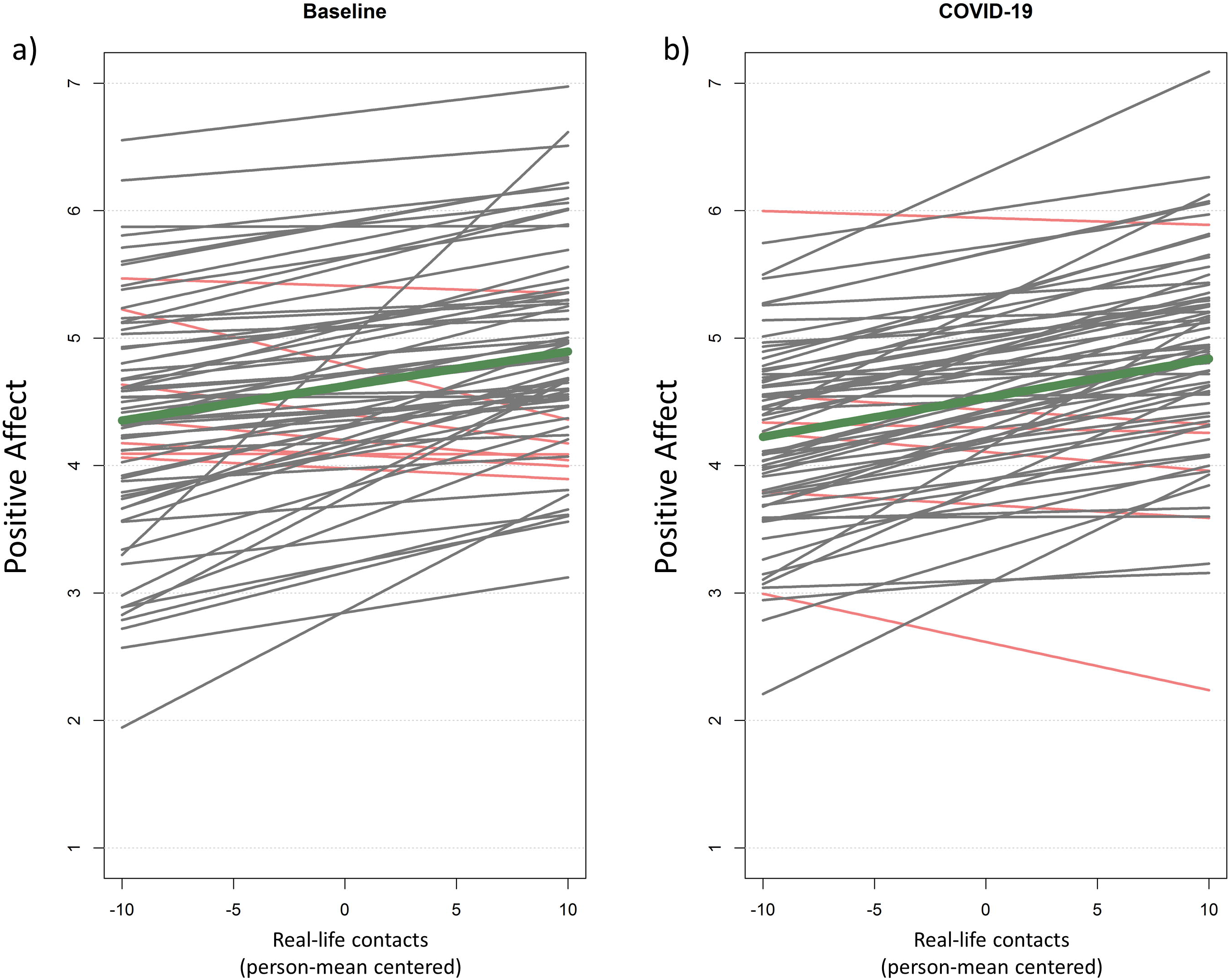
Individual associations of real-life contacts and positive affect during baseline (a) and during COVID-19 (b). Real-life contacts represent person-mean centered social contacts within the last two hours. Differences from zero indicate an increased / decreased amount of social contacts compared to the person-mean. Gray and red lines reflect positive and negative slope values, respectively. Thick line in dark green reflects the association for the whole group. Notably, decreased overall positive affect during the COVID-19 pandemic was not fully explained by social contact reductions.

### Amygdala volume, social contacts, and affective state (Models-II and III)

Left but not right amygdala volume moderated the relationship between the number of social contacts, time point and positive affect (P=0.008, Table 1; P=0.675, Supplementary Table 4a, respectively).

**Table 1:**
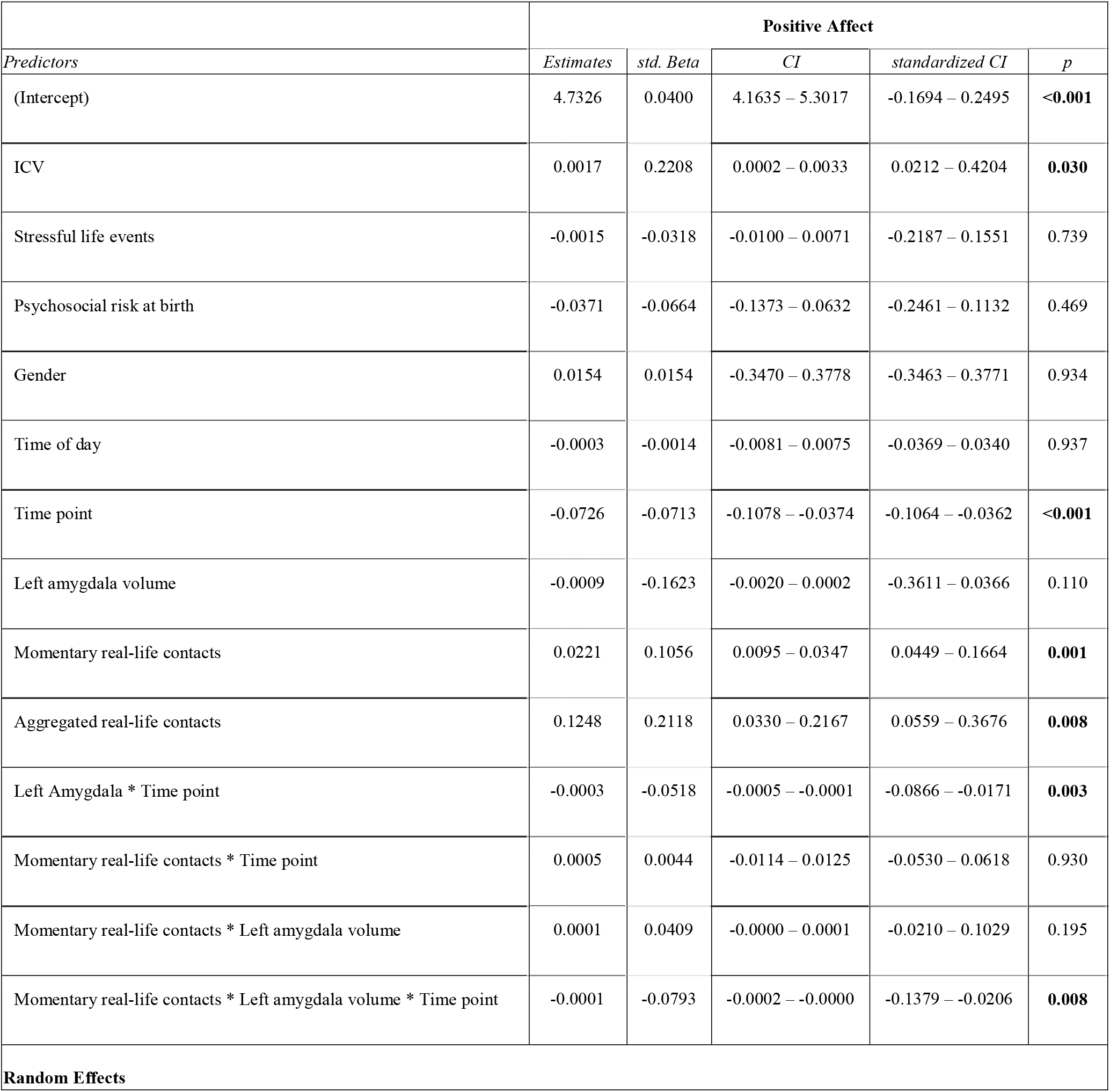
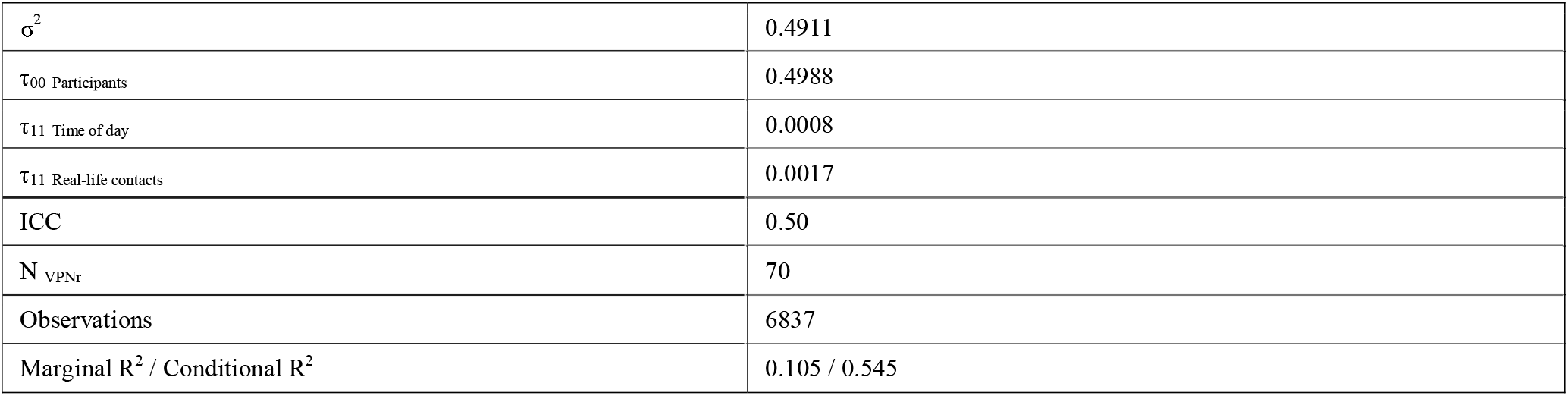
Mixed model results for social contacts, left amygdala volume, and positive affective states across both time points

| *Predictors* | **Positive Affect** |  |  |  |  |
| --- | --- | --- | --- | --- | --- |
|  | *Estimates* | *std. Beta* | *CI* | *standardized CI* | *p* |
| (Intercept) | 4.7326 | 0.0400 | 4.1635 – 5.3017 | -0.1694 – 0.2495 | **<0.001** |
| ICV | 0.0017 | 0.2208 | 0.0002 – 0.0033 | 0.0212 – 0.4204 | **0.030** |
| Stressful life events | -0.0015 | -0.0318 | -0.0100 – 0.0071 | -0.2187 – 0.1551 | 0.739 |
| Psychosocial risk at birth | -0.0371 | -0.0664 | -0.1373 – 0.0632 | -0.2461 – 0.1132 | 0.469 |
| Gender | 0.0154 | 0.0154 | -0.3470 – 0.3778 | -0.3463 – 0.3771 | 0.934 |
| Time of day | -0.0003 | -0.0014 | -0.0081 – 0.0075 | -0.0369 – 0.0340 | 0.937 |
| Time point | -0.0726 | -0.0713 | -0.1078 – -0.0374 | -0.1064 – -0.0362 | **<0.001** |
| Left amygdala volume | -0.0009 | -0.1623 | -0.0020 – 0.0002 | -0.3611 – 0.0366 | 0.110 |
| Momentary real-life contacts | 0.0221 | 0.1056 | 0.0095 – 0.0347 | 0.0449 – 0.1664 | **0.001** |
| Aggregated real-life contacts | 0.1248 | 0.2118 | 0.0330 – 0.2167 | 0.0559 – 0.3676 | **0.008** |
| Left Amygdala * Time point | -0.0003 | -0.0518 | -0.0005 – -0.0001 | -0.0866 – -0.0171 | **0.003** |
| Momentary real-life contacts * Time point | 0.0005 | 0.0044 | -0.0114 – 0.0125 | -0.0530 – 0.0618 | 0.930 |
| Momentary real-life contacts * Left amygdala volume | 0.0001 | 0.0409 | -0.0000 – 0.0001 | -0.0210 – 0.1029 | 0.195 |
| Momentary real-life contacts * Left amygdala volume * Time point | -0.0001 | -0.0793 | -0.0002 – -0.0000 | -0.1379 – -0.0206 | **0.008** |
| **Random Effects** |  |  |  |  |  |

|  |  |
| --- | --- |
| $\sigma^2$ | 0.4911 |
| $\tau_{00}$ Participants | 0.4988 |
| $\tau_{11}$ Time of day | 0.0008 |
| $\tau_{11}$ Real-life contacts | 0.0017 |
| ICC | 0.50 |
| $N_{VPNr}$ | 70 |
| Observations | 6837 |
| Marginal $R^2$ / Conditional $R^2$ | 0.105 / 0.545 |

Subsequent simple slope analyses and Johnson-Neyman plots (Figure 2a&2b) for the left amygdala showed a significant positive association (i.e., a positive slope) between real-life contacts and positive affect depending on the amygdala size, indicating an affective gain from social interactions only in those with larger amygdala volumes. In contrast, the opposite relationship was present during the lockdown, with an affective gain only in those with average to smaller amygdala volumes.

**Figure 2:**
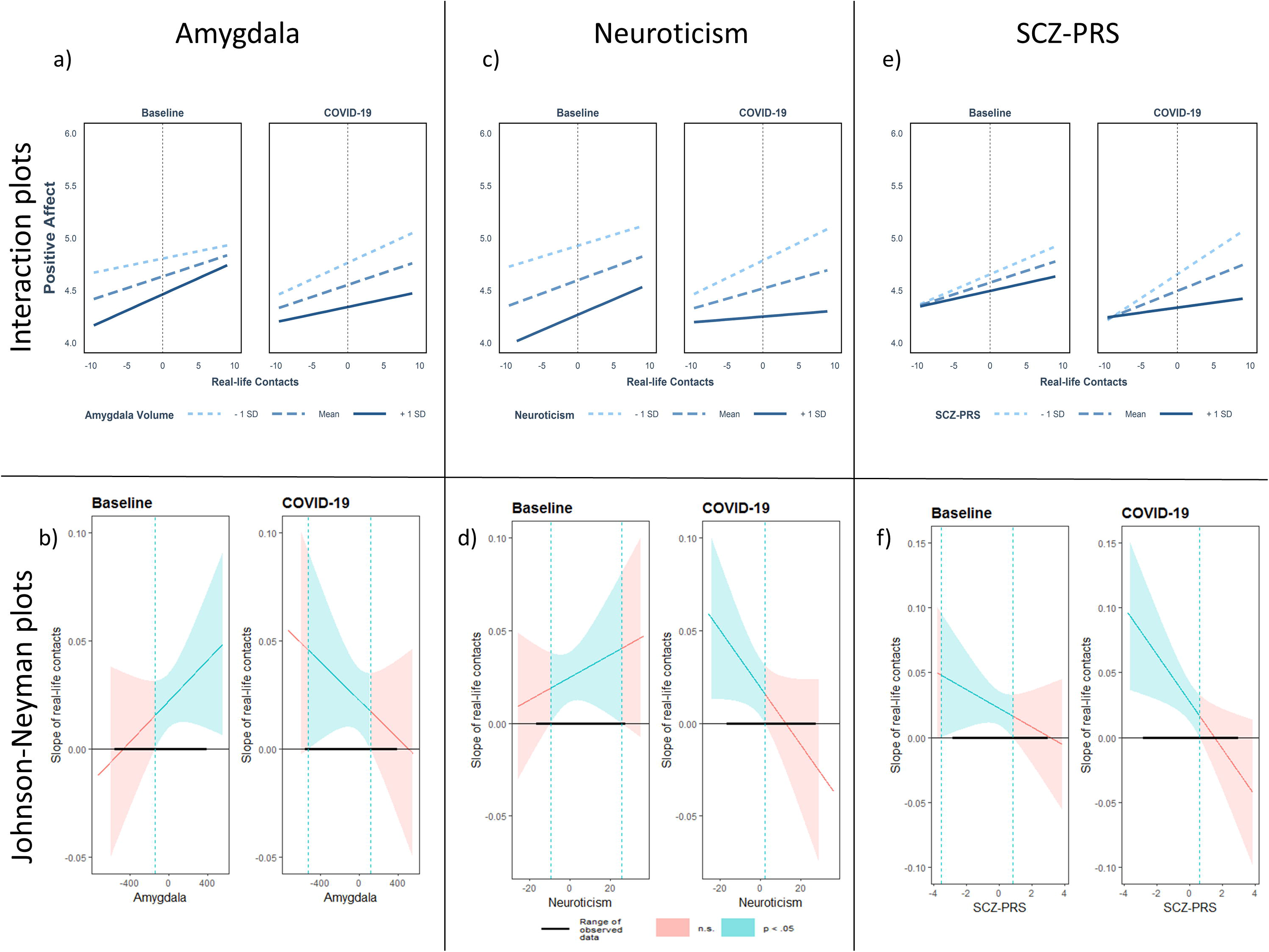
Interaction- and Johnson-Neyman plots for all significant three-way interactions with the quantity of real-life contacts. Top: Plots depicting the interaction between real-life contacts, time point and amygdala (a), neuroticism (c), and polygenic risk for schizophrenia (SCZ-PRS, e) on positive affect. Real-life contacts represent person-mean centered social contacts within the last two hours. Differences from zero indicate an increased / decreased amount of social contacts compared to the person-mean. Bottom: Johnson-Neyman plots for the significant three-way interactions with amygdala (b), neuroticism (d), and SCZ-PRS (f). Johnson-Neyman plots indicate the range of observed values of a moderator, for which the association (i.e. ‘slope of real-life contacts’) between real-life contacts and positive affect is significant (p<0.05).

### Neuroticism, social contacts and affective state (Model-IV)

The results revealed significant three-way interactions between neuroticism, time point and the quantity of real-life contacts (P=0.003, Table 2). Subsequent analyses for real-life contacts indicated an affective benefit from social interactions irrespective of trait neuroticism before the pandemic. However, during the COVID-19 assessment, the beneficial effect of real-life contacts on positive affect was only significant in those with low neuroticism scores, whereas those with high neurotic traits showed no affective benefit (Figure 2c&2d).

**Table 2:**
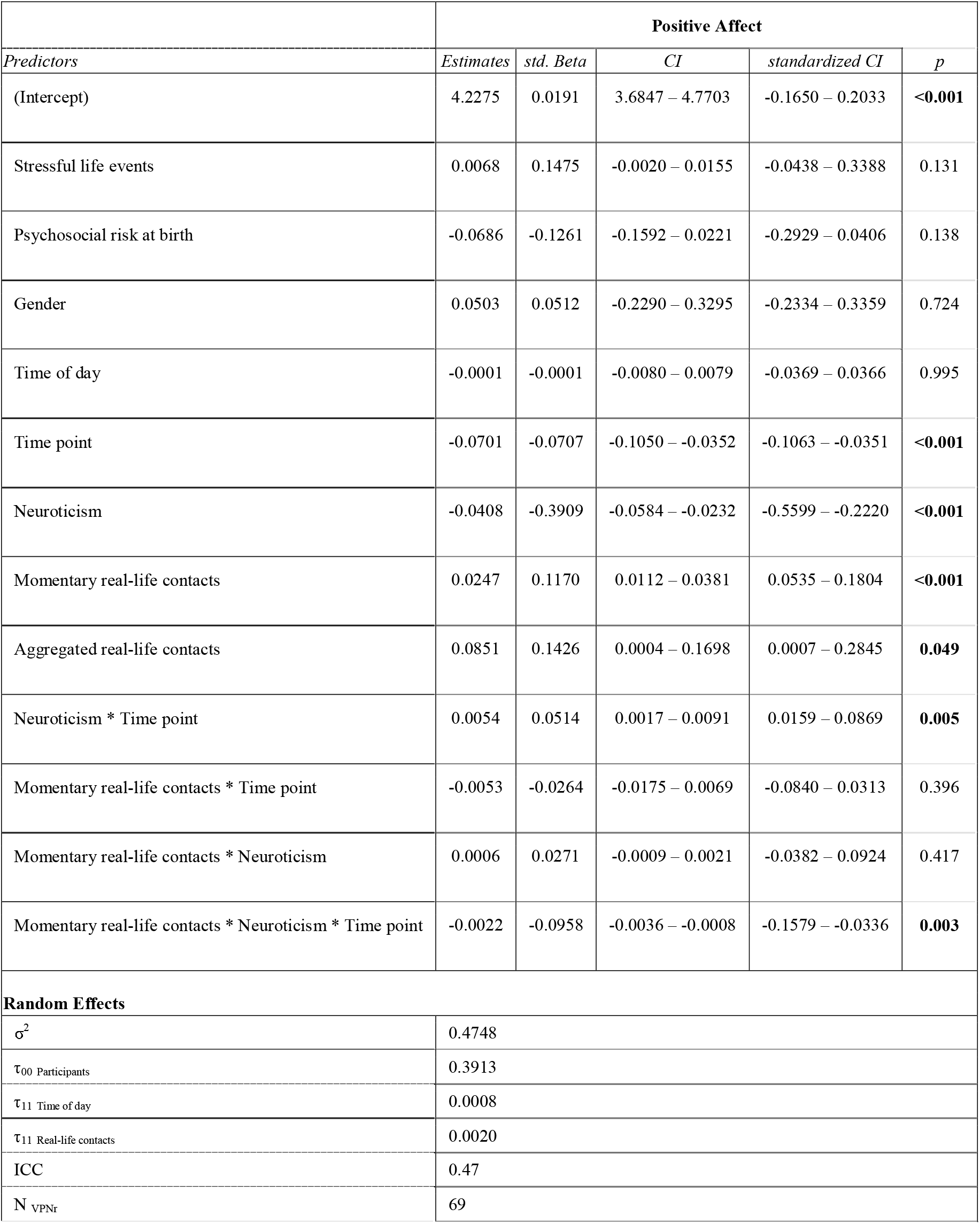
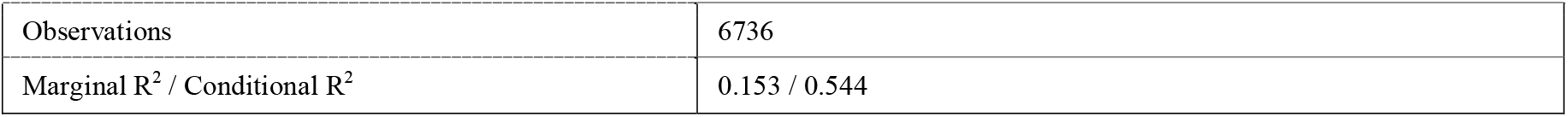
Mixed model results for social contacts, neuroticism, and positive affective states across both time points

### Polygenic risk scores, social contacts, and affective state (Model-V)

Finally, there was a three-way interaction for SCZ-PRS, quantity of social contacts, and time point on positive affect (real-life contacts: P = 0.036, Table 3). Subsequent analyses revealed that those with low to moderate SCZ-PRS showed an affective benefit from social interactions before COVID-19 (Figure 2e&2f). This relationship between real-life contacts and positive affect even increased during COVID-19 in participants with low to moderate SCZ-PRS.

**Table 3:**
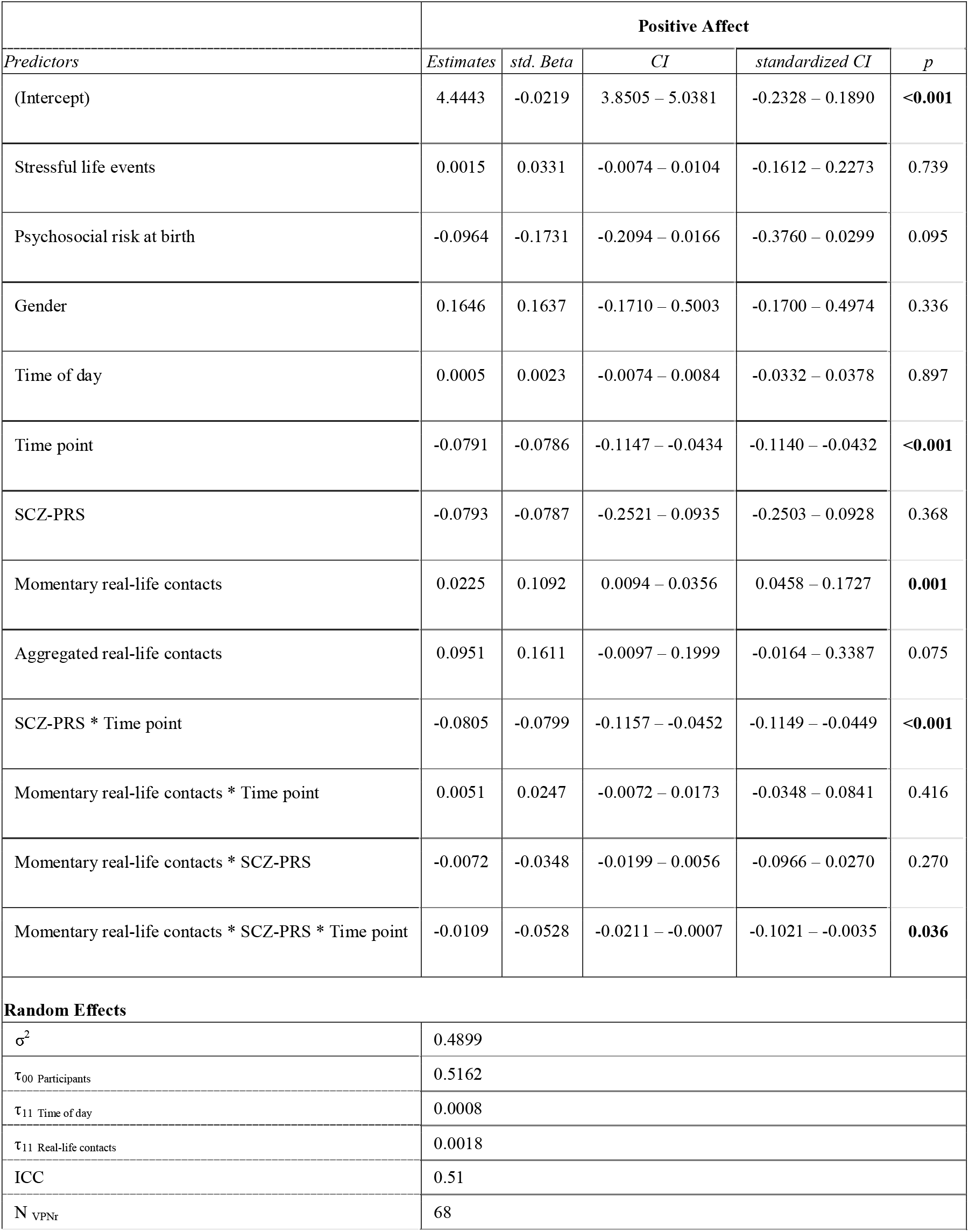
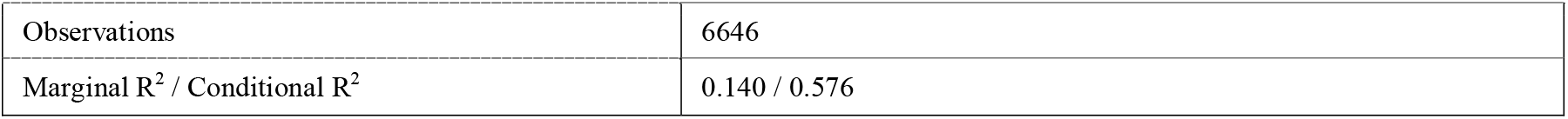
Mixed model results for social contacts, SCZ-PRS, and positive affective states across both time points

| *Predictors* | **Positive Affect** |  |  |  |  |
| --- | --- | --- | --- | --- | --- |
|  | *Estimates* | *std. Beta* | *CI* | *standardized CI* | *p* |
| (Intercept) | 4.4443 | -0.0219 | 3.8505 – 5.0381 | -0.2328 – 0.1890 | **<0.001** |
| Stressful life events | 0.0015 | 0.0331 | -0.0074 – 0.0104 | -0.1612 – 0.2273 | 0.739 |
| Psychosocial risk at birth | -0.0964 | -0.1731 | -0.2094 – 0.0166 | -0.3760 – 0.0299 | 0.095 |
| Gender | 0.1646 | 0.1637 | -0.1710 – 0.5003 | -0.1700 – 0.4974 | 0.336 |
| Time of day | 0.0005 | 0.0023 | -0.0074 – 0.0084 | -0.0332 – 0.0378 | 0.897 |
| Time point | -0.0791 | -0.0786 | -0.1147 – -0.0434 | -0.1140 – -0.0432 | **<0.001** |
| SCZ-PRS | -0.0793 | -0.0787 | -0.2521 – 0.0935 | -0.2503 – 0.0928 | 0.368 |
| Momentary real-life contacts | 0.0225 | 0.1092 | 0.0094 – 0.0356 | 0.0458 – 0.1727 | **0.001** |
| Aggregated real-life contacts | 0.0951 | 0.1611 | -0.0097 – 0.1999 | -0.0164 – 0.3387 | 0.075 |
| SCZ-PRS * Time point | -0.0805 | -0.0799 | -0.1157 – -0.0452 | -0.1149 – -0.0449 | **<0.001** |
| Momentary real-life contacts * Time point | 0.0051 | 0.0247 | -0.0072 – 0.0173 | -0.0348 – 0.0841 | 0.416 |
| Momentary real-life contacts * SCZ-PRS | -0.0072 | -0.0348 | -0.0199 – 0.0056 | -0.0966 – 0.0270 | 0.270 |
| Momentary real-life contacts * SCZ-PRS * Time point | -0.0109 | -0.0528 | -0.0211 – -0.0007 | -0.1021 – -0.0035 | **0.036** |
| **Random Effects** |  |  |  |  |  |
| $\sigma^2$ | 0.4899 | | | | |
| $\tau_{00}$ Participants | 0.5162 | | | | |
| $\tau_{11}$ Time of day | 0.0008 | | | | |
| $\tau_{11}$ Real-life contacts | 0.0018 | | | | |
| ICC | 0.51 |  |  |  |  |
| N VPnr | 68 |  |  |  |  |

|  |  |
| --- | --- |
| Observations | 6646 |
| Marginal $R^2$ / Conditional $R^2$ | 0.140 / 0.576 |

## Discussion

To the best of our knowledge, this is the first longitudinal study to investigate the social affective benefit in a real-time, real-life setting prior to and during the lockdown. Our results demonstrate that both the quantity and the quality of social interactions were significantly associated with positive affect, irrespective of the assessed time point (Model I). In addition, we demonstrate for the first time that amygdala volume (Model II-III), neuroticism (Model IV) and SCZ-PRS (Model V) moderated the relationship between the number of social contacts and positive affect differently during normal times as compared to the lockdown.

In line with previous findings (11), the number and quality of real-life social interactions exerted mood-uplifting effects, both before and during the lockdown, indicating an overall protective role of frequent social contacts on well-being. Thereby, we critically expand those findings by showing that this relationship is stable even under social contact restrictions.

Moreover, we demonstrate that the beneficial impact of frequent social interactions on well-being is determined by an individual’s neurobiobehavioral profile. As such, left amygdala volume (Model-II) moderated the social affective gain differently before and during Covid-19. This finding is in accordance with the current consensus embedding the amygdala in stress adaptation (51) and a left lateralized sustained involvement in emotional contexts (52–54). Here, we extend these previous findings by suggesting that one possible mechanism underlying stress adaptation pertains to left amygdala-dependent modulation of the beneficial effect of social interactions. In line with previous reports linking larger amygdala volumes to increased social network sizes and to higher levels of social support (15, 17), only participants with larger amygdala volumes showed a social affective benefit before the lockdown, indicating that these individuals might more actively seek for social interactions to improve positive emotionality. However, this pattern changed under social contact restrictions, with only participants with average to low left amygdala volumes benefiting from more real-life contacts during the lockdown phase of the COVID-19 pandemic. In this regard, it seems that individuals with a larger left amygdala volume may no longer offset their lower positive affect by increasing social contacts under stress when compared to normal times. This provides a mechanistic understanding of how a larger amygdala volume can be considered a risk marker which only manifests under adversity.

Similarly to those with a larger amygdala volume, participants with higher neurotic traits benefit from more social contacts (Model IV) before Covid-19, thus may catching up on their affective backlog (55). Interestingly, this pattern is reversed during the lockdown, indicating an expected mood-lifting effect of social interactions only in persons with low to moderate levels of neuroticism. This is consistent with the idea that individuals with high neurotic traits experience uneasy times as more negative and aversive and are less equipped with adequate coping strategies, such as seeking for social support (30). We speculate that participants with higher neurotic traits show more interactions focused on COVID-19 related information. That is in line with previous studies during the COVID-19 pandemic reporting higher neurotic traits to be associated with decreased overall well-being during the COVID-19 pandemic (20). Therefore, our results suggest that high levels of neuroticism might be particularly maladaptive on social affective gain during times of crisis, while their impact on the social affective benefit might be less manifest during normal times.

Finally, we found that the genetic risk for schizophrenia moderated the relationship between social contacts and positive affect (Model-V). As expected and in line with previous findings (32), we found that only those with lower genetic risk for schizophrenia showed an affective gain from social interactions Notably, while this has been previously shown during normal times (32), our results show that this is aggravated under social contact restrictions. Therefore, enhancing coping strategies in those at risk might prove beneficial to prevent the onset of psychotic, particularly negative, symptoms in individuals at risk. However, this analysis has to be considered exploratory, given that it fell short of significance after multiple comparison correction.

Some limitations of the present study need to be addressed. Since the study started during social contact restrictions, only a quarter of our MARS participants were able to take part in this follow-up measurement. However, the extremely high compliance rate of 87% guaranteed enough power and the sample was not systematically biased regarding demographics and psychosocial risk factors when compared to the dropout sample. Moreover, since the COVID-19 infection rates were relatively low in Germany, an underestimation of the physical and mental health consequences within our sample cannot be ruled out.

Taken together, our findings highlight the protective role of social interactions on well-being both before and during the pandemic lockdown. Compared to normal times, individuals with smaller amygdala, low levels of neuroticisms and low risk for schizophrenia demonstrated an increased social affective benefit during social contact restrictions. In this respect, our findings demonstrate critical determinants of social affective gain, which might act as targets for future interventions.

## Acknowledgments

The authors gratefully thank Manfred Laucht (1946-2020), who was one of the founders of the Mannheim Study of Children at Risk and who continuously acted as an inspiring and supporting mentor giving impulses for innovative research projects.

This work was gratefully supported by the German Research Foundation (grant number DFG HO 5674/2-1, GRK2350/1) and the Ministry of Science, Research and the Arts of the State of Baden-Württemberg, Germany (Special support program SARS CoV-2 pandemic) to N.H. T.B. gratefully acknowledges grant support by the German Federal Ministry of Education and Research (01EE1408E ESCAlife; FKZ 01GL1741[X] ADOPT; 01EE1406C Verbund AERIAL; 01EE1409C Verbund ASD-Net; 01GL1747C STAR; 01GL1745B IMAC-Mind), by the German Research Foundation (TRR 265/1), by the Innovative Medicines Initiative Joint Undertaking (IMI JU FP7 115300 EU-AIMS; grant 777394 EU-AIMS-2-TRIALS) and the European Union – H2020 (Eat2beNICE, grant 728018; PRIME, grant 847879).

A.M.L. and H.T. acknowledge grant support by the German Research Foundation (DFG, Research Training Group GRK2350/1 project B02, Collaborative Research Center SFB 1158 project B09, Collaborative Research Center TRR 265 project S02, grant ME 1591/4-1), German Federal Ministry of Education and Research (BMBF, grants 01EF1803A, 01ZX1314G, 01GQ1003B), European Union’s Seventh Framework Programme (FP7, grants 602450, 602805, 115300, HEALTH-F2-2010-241909, Horizon2020 CANDY grant 847818 and Eat2beNICE grant 728018), Innovative Medicines Initiative Joint Undertaking (IMI, grant 115008, PRISM grant 115916, EU-AIMS grant 115300, and AIMS-2-TRIALS grant 777394) and Ministry of Science, Research and the Arts of the State of Baden-Wuerttemberg, Germany (MWK, grant 42-04HV.MED(16)/16/1).

Additionally, this work was supported by the German Federal Ministry of Education and Research (BMBF) through ERA-NET NEURON, “SynSchiz - Linking synaptic dysfunction to disease mechanisms in schizophrenia - a multilevel investigation” [01EW1810] and through ERA-NET NEURON “Impact of Early life MetaBolic and psychosocial strEss on susceptibility to mental Disorders; from converging epigenetic signatures to novel targets for therapeutic intervention” [01EW1904]. The funding sources had no role in study design, collection, analysis, and interpretation of data, in the writing of the report, and in the decision to submit the article for publication. The authors thank the MARS participants for their continuous participation in the assessments. The authors further thank Sibylle Heinzel, Rafaela Gehr, Christin Loebel and Cäcilia Pracht for supporting the assessments.

## References

1. Qiu J, Shen B, Zhao M, Wang Z, Xie B, Xu Y (2020): A nationwide survey of psychological distress among Chinese people in the COVID-19 epidemic: implications and policy recommendations. Gen Psychiatr. 33:e100213.

2. Wang ZH, Yang HL, Yang YQ, Liu D, Li ZH, Zhang XR, et al. (2020): Prevalence of anxiety and depression symptom, and the demands for psychological knowledge and interventions in college students during COVID-19 epidemic: A large cross-sectional study. J Affect Disord. 275:188–193.

3. Rafnsson SB, Shankar A, Steptoe A (2015): Longitudinal Influences of Social Network Characteristics on Subjective Well-Being of Older Adults: Findings From the ELSA Study. J Aging Health. 27:919–934.

4. Zhang Z, Zhang J, Zhao N, Yang Y (2019): Social Network Size and Subjective Well-Being: The Mediating Role of Future Time Perspective Among Community-Dwelling Retirees. Front Psychol. 10:2590.

5. Smyth JM, Zawadzki MJ, Santuzzi AM, Filipkowski KB (2014): Examining the effects of perceived social support on momentary mood and symptom reports in asthma and arthritis patients. Psychology & health. 29:813–831.

6. Bloom JR, Stewart SL, Johnston M, Banks P, Fobair P (2001): Sources of support and the physical and mental well-being of young women with breast cancer. Social Science & Medicine. 53:1513–1524.

7. Cacioppo JT, Hughes ME, Waite LJ, Hawkley LC, Thisted RA (2006): Loneliness as a specific risk factor for depressive symptoms: cross-sectional and longitudinal analyses. Psychol Aging. 21:140–151.

8. Pantell M, Rehkopf D, Jutte D, Syme SL, Balmes J, Adler N (2013): Social isolation: a predictor of mortality comparable to traditional clinical risk factors. Am J Public Health. 103:2056–2062.

9. Holt-Lunstad J, Smith TB, Baker M, Harris T, Stephenson D (2015): Loneliness and social isolation as risk factors for mortality: a meta-analytic review. Perspect Psychol Sci. 10:227–237.

10. Rico-Uribe LA, Caballero FF, Martin-Maria N, Cabello M, Ayuso-Mateos JL, Miret M (2018): Association of loneliness with all-cause mortality: A meta-analysis. PLoS One. 13:e0190033.

11. Bernstein MJ, Zawadzki MJ, Juth V, Benfield JA, Smyth JM (2017): Social interactions in daily life. Journal of Social and Personal Relationships. 35:372–394.

12. Elmer T, Mepham K, Stadtfeld C (2020): Students under lockdown: Comparisons of students’ social networks and mental health before and during the COVID-19 crisis in Switzerland. PLoS One. 15:e0236337.

13. Bickart KC, Dickerson BC, Barrett LF (2014): The amygdala as a hub in brain networks that support social life. Neuropsychologia. 63:235–248.

14. Holz NE, Tost H, Meyer-Lindenberg A (2019): Resilience and the brain: a key role for regulatory circuits linked to social stress and support. Mol Psychiatry. 25:379–396.

15. Bickart KC, Wright CI, Dautoff RJ, Dickerson BC, Barrett LF (2011): Amygdala volume and social network size in humans. Nat Neurosci. 14:163–164.

16. Kanai R, Bahrami B, Roylance R, Rees G (2012): Online social network size is reflected in human brain structure. Proc Biol Sci. 279:1327–1334.

17. Sato W, Kochiyama T, Kubota Y, Uono S, Sawada R, Yoshimura S, et al. (2016): The association between perceived social support and amygdala structure. Neuropsychologia. 85:237–244.

18. Duzel S, Drewelies J, Gerstorf D, Demuth I, Steinhagen-Thiessen E, Lindenberger U, et al. (2019): Structural Brain Correlates of Loneliness among Older Adults. Sci Rep. 9:13569.

19. Kalish Y, Robins G (2006): Psychological predispositions and network structure: The relationship between individual predispositions, structural holes and network closure. Social Networks. 28:56–84.

20. Shokrkon A, Nicoladis E (2021): How personality traits of neuroticism and extroversion predict the effects of the COVID-19 on the mental health of Canadians. PLoS One. 16:e0251097.

21. van Os J, Kenis G, Rutten BP (2010): The environment and schizophrenia. Nature. 468:203–212.

22. Tost H, Champagne FA, Meyer-Lindenberg A (2015): Environmental influence in the brain, human welfare and mental health. Nat Neurosci. 18:1421–1431.

23. Robustelli BL, Newberry RE, Whisman MA, Mittal VA (2017): Social relationships in young adults at ultra high risk for psychosis. Psychiatry Res. 247:345–351.

24. Goldberg RW, Rollings AL, Lehman AF (2003): Social Network Correlates among People with Psychiatric Disabilities. Psychiatric Rehabilitation Journal. 26:393–402.

25. Kraaijenvanger EJ, Pollok TM, Monninger M, Kaiser A, Brandeis D, Banaschewski T, et al. (2020): Impact of early life adversities on human brain functioning: A coordinate-based meta-analysis. Neurosci Biobehav Rev. 113:62–76.

26. Sublette ME, Galfalvy HC, Oquendo MA, Bart CP, Schneck N, Arango V, et al. (2016): Relationship of recent stress to amygdala volume in depressed and healthy adults. J Affect Disord. 203:136–142.

27. Roozendaal B, McEwen BS, Chattarji S (2009): Stress, memory and the amygdala. Nat Rev Neurosci. 10:423–433.

28. Quinlan EB, Cattrell A, Jia T, Artiges E, Banaschewski T, Barker G, et al. (2017): Psychosocial Stress and Brain Function in Adolescent Psychopathology. Am J Psychiatry. 174:785–794.

29. Farmer A, Redman K, Harris T, Mahmood A, Sadler S, Pickering A, et al. (2002): Neuroticism, extraversion, life eventseve and depression. British Journal of Psychiatry. 181:118 – 122.

30. Carver CS, Connor-Smith J (2010): Personality and coping. Annu Rev Psychol. 61:679–704.

31. Castellanos-Ryan N, Briere FN, O’Leary-Barrett M, Banaschewski T, Bokde A, Bromberg U, et al. (2016): The structure of psychopathology in adolescence and its common personality and cognitive correlates. J Abnorm Psychol. 125:1039–1052.

32. Myin-Germeys I, van Os J, Schwartz JE, Stone AA, Delespaul PA (2001): Emotional Reactivity to Daily Life Stress in Psychosis. Arch Gen Psychiatry 58:1137–1144.

33. Morey RA, Gold AL, LaBar KS, Beall SK, Brown VM, Haswell CC, et al. (2012): Amygdala Volume Changes in Posttraumatic Stress Disorder in a Large Case-Controlled Veterans Group. Arch Gen Psychiatry 69:1169–1178.

34. Gunthert KC, Cohen LH, Armeli S (1999): The Role of Neuroticism in Daily Stress and Coping. Journal of Personality and Social Psychology. 77:1087–1100.

35. Laucht M, Esser G, Baving L, Gerhold M, Hoesch I, Ihle W, et al. (2000): Behavioral Sequelae of Perinatal Insults and Early Family Adversity at 8 Years of Age. J Am Acad Child Adolesc Psychiatry. 39:1229–1237.

36. Krohne HW, Egloff B, Kohlmann C-W, Tausch A (1996): Untersuchungen mit einer deutschen Version der “Positive and Negative Affect Schedule” (PANAS). Diagnostica. 42:139–159.

37. Hohn P, Menne-Lothmann C, Peeters F, Nicolson NA, Jacobs N, Derom C, et al. (2013): Moment-to-moment transfer of positive emotions in daily life predicts future course of depression in both general population and patient samples. PLoS One. 8:e75655.

38. Myin-Germeys I, Peeters F, Havermans R, Nicolson NA, deVries MW, Delespaul P, et al. (2003): Emotional reactivity to daily life stress in psychosis and affective disorders: an experience sampling study. Acta Psychiatr Scand. 107:124–131.

39. van de Leemput IA, Wichers M, Cramer AO, Borsboom D, Tuerlinckx F, Kuppens P, et al. (2013): Critical slowing down as early warning for the onset and termination of depression. Proc Natl Acad Sci U S A. 111:87–92.

40. Wichers M, Peeters F, Geschwind N, Jacobs N, Simons CJ, Derom C, et al. (2010): Unveiling patterns of affective responses in daily life may improve outcome prediction in depression: a momentary assessment study. J Affect Disord. 124:191–195.

41. Shrout PE, Lane SP (2012): Psychometrics.

42. Fischl B (2012): FreeSurfer. Neuroimage. 62:774–781.

43. Borkenau P, Ostendorf F (2008): NEO-FFI: NEO-Fünf-Faktoren Inventar nach Costa und McCrae. Hogrefe.

44. Chang CC, Chow CC, Tellier LC, Vattikuti S, Purcell SM, Lee JJ (2015): Second-generation PLINK: rising to the challenge of larger and richer datasets. GigaScience. 4.

45. Schizophrenia Working Group of the Psychiatric Genomics C (2014): Biological insights from 108 schizophrenia-associated genetic loci. Nature. 511:421–427.

46. Choi SW, O’Reilly PF (2019): PRSice-2: Polygenic Risk Score software for biobank-scale data. GigaScience. 8.

47. Rutter M, Quinton D (1977): Psychiatric disorder: ecological factors and concepts of causation. In: McGurk M, editor. Ecological factors in human development: Amsterdam: Noord-Holland, pp 173–187.

48. Maier-Diewald W (1983): Die Münchner Ereignisliste: MEL Max-Planck-Institut für Psychiatrie.

49. Bates D, Mächler M, Bolker B, Walker S (2015): Fitting Linear Mixed-Effects Models Using lme4. 2015. 67:48.

50. Kuznetsova A, Brockhoff PB, Christensen RHB (2017): lmerTest Package: Tests in Linear Mixed Effects Models. Journal of Statistical Software. 82.

51. McEwen BS, Gianaros PJ (2010): Central role of the brain in stress and adaptation: links to socioeconomic status, health, and disease. Ann N Y Acad Sci. 1186:190–222.

52. Baas D, Aleman A, Kahn RS (2004): Lateralization of amygdala activation: a systematic review of functional neuroimaging studies. Brain Research Reviews. 45:96–103.

53. Wager TD, Phan KL, Liberzon I, Taylor SF (2003): Valence, gender, and lateralization of functional brain anatomy in emotion: a meta-analysis of findings from neuroimaging. NeuroImage. 19:513–531.

54. Sergerie K, Chochol C, Armony JL (2008): The role of the amygdala in emotional processing: a quantitative meta-analysis of functional neuroimaging studies. Neurosci Biobehav Rev. 32:811–830.

55. Steger MF, Kashdan TB (2009): Depression and Everyday Social Activity, Belonging, and Well-Being. J Couns Psychol. 56:289–300.

