## Supplementary figures and images for "Real-time individual benefit from social interactions before and during the lockdown: The crucial role of personality, neurobiology and genes"

### Supplemental Figure 1

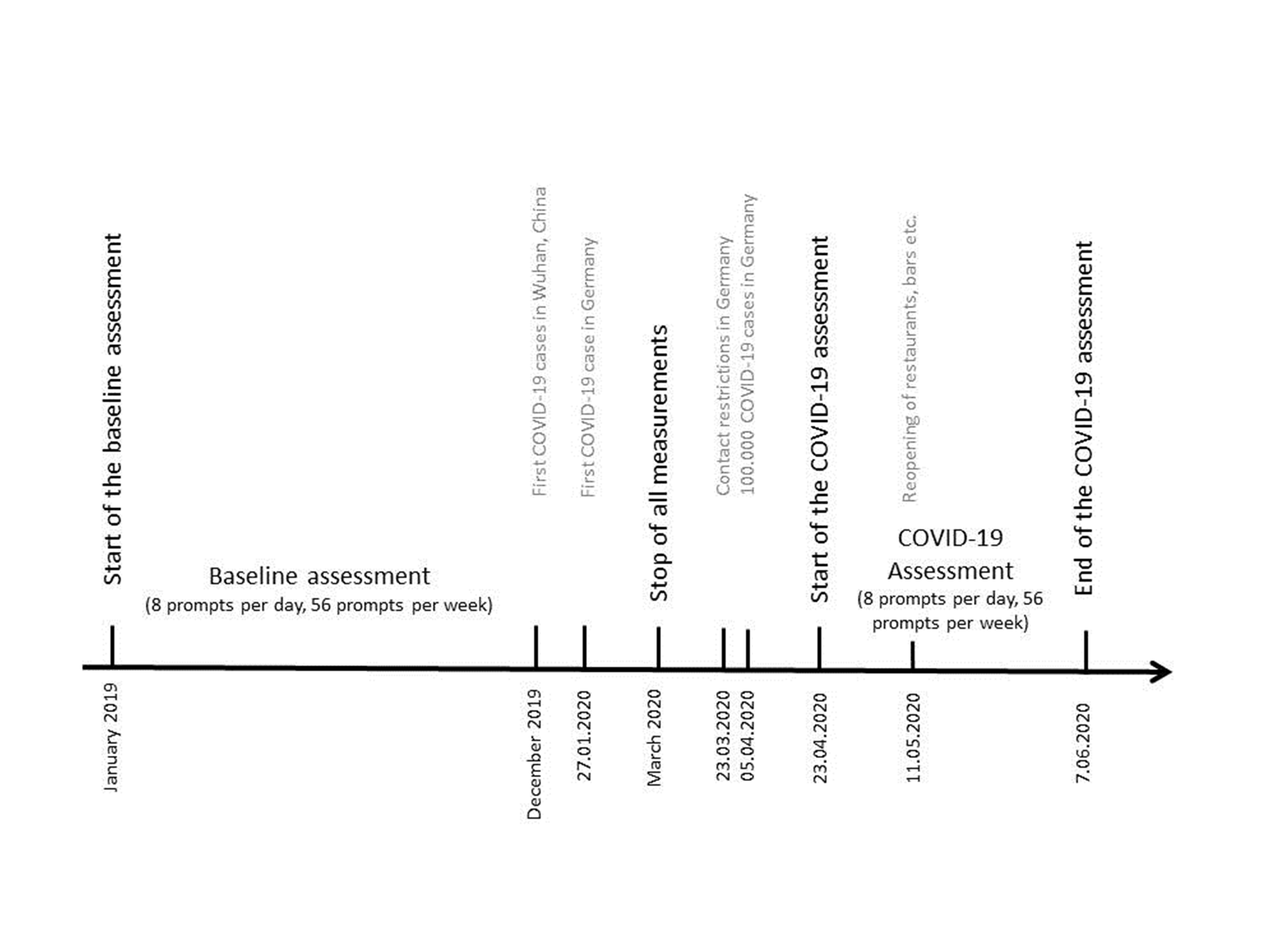
