## Supplemental Tables for "Real-time individual benefit from social interactions before and during the lockdown: The crucial role of personality, neurobiology and genes"

### Supplementary Table 1.

Descriptive data

| **Descriptive data (n = 70)** | | | |
| --- | --- | --- | --- |
|  | *Mean* | *SD* | *Range* |
| Positive affect | 4.60 | 1 | 1 - 7 |
| Negative affect | 1.47 | 0.78 | 1 - 5.22 |
| COVID-19 stress burden | 7.93 | 4.84 | 2 - 18 |
| Stressful life events | 58.11 | 21.77 | 28 - 129 |
| Psychosocial Risk factors at birth | 1.87 | 1.8 | 0 - 7 |
| Neuroticism | 18.28 | 9.06 | 2 - 39 |
| SCZ-PRS | -0.0015 | 0.0001 | -0.0017 - -0.0012 |
| Left Amygdala Volume (mm³) | 1633.7 | 179.87 | 1089 - 2017 |
| Right Amygdala Volume (mm³) | 1727.1 | 204.8 | 1301 - 2176 |
| Estimated Total Intracranial Volume (mm³) | 1503564 | 128345 | 1257021 - 1932126 |
| Age (years) | 33.37 | 0.57 | 32.25 - 34.25 |

### Supplementary Table 2a.

Mixed model results for social contacts and positive affective states across both time points

|  | **Positive Affect** | | | | |
| --- | --- | --- | --- | --- | --- |
| *Predictors* | *Estimates* | *std. Beta* | *CI* | *standardized CI* | *p* |
| (Intercept) | 4.7108 | 0.0054 | 4.1600 – 5.2616 | -0.1959 – 0.2067 | <0.001 |
| Stressful life events | -0.0017 | -0.0366 | -0.0102 – 0.0069 | -0.2226 – 0.1495 | 0.700 |
| Psychosocial risk at birth | -0.0433 | -0.0776 | -0.1451 – 0.0586 | -0.2602 – 0.1050 | 0.405 |
| Gender | 0.1002 | 0.0999 | -0.2160 – 0.4163 | -0.2156 – 0.4155 | 0.535 |
| Time of day | -0.0002 | -0.0011 | -0.0081 – 0.0076 | -0.0366 – 0.0343 | 0.951 |
| Time point | -0.0738 | -0.0737 | -0.1090 – -0.0386 | -0.1088 – -0.0385 | **<0.001** |
| Momentary real-life contacts | 0.0215 | 0.1035 | 0.0087 – 0.0343 | 0.0420 – 0.1651 | **0.001** |
| Aggregated real-life contacts | 0.1343 | 0.2278 | 0.0424 – 0.2262 | 0.0719 – 0.3837 | **0.004** |
| Momentary real-life contacts * Time point | 0.0010 | 0.0047 | -0.0109 – 0.0129 | -0.0525 – 0.0620 | 0.871 |
| **Random Effects** | | | | | |
| σ^2^ | 0.4917 | | | | |
| τ_00_ _Participants_ | 0.5164 | | | | |
| τ_11_ _Time of day_ | 0.0008 | | | | |
| τ_11_ _Real-life contacts_ | 0.0018 | | | | |
| ICC | 0.50 | | | | |
| N _VPNr_ | 70 | | | | |
| Observations | 6837 | | | | |
| Marginal R^2^ / Conditional R^2^ | 0.074 / 0.540 | | | | |

### Supplementary Table 2b.

Mixed model results for social contacts and positive affective states across both time points

|  | **Positive Affect** | | | | |
| --- | --- | --- | --- | --- | --- |
| *Predictors* | *Estimates* | *std. Beta* | *CI* | *standardized CI* | *p* |
| (Intercept) | 4.7481 | -0.0343 | 4.2236 – 5.2726 | -0.2351 – 0.1665 | <0.001 |
| Stressful life events | -0.0018 | -0.0424 | -0.0099 – 0.0062 | -0.2283 – 0.1435 | 0.655 |
| Psychosocial risk at birth | -0.0360 | -0.0668 | -0.1329 – 0.0610 | -0.2467 – 0.1132 | 0.467 |
| Gender | 0.1908 | 0.1981 | -0.1096 – 0.4913 | -0.1138 – 0.5100 | 0.213 |
| Time of day | -0.0030 | -0.0138 | -0.0102 – 0.0042 | -0.0473 – 0.0196 | 0.417 |
| Time point | -0.0963 | -0.1031 | -0.1319 – -0.0607 | -0.1407 – -0.0655 | **<0.001** |
| Momentary real-life contacts | 0.0132 | 0.0693 | 0.0042 – 0.0222 | 0.0222 – 0.1165 | **0.004** |
| Quality of most important interaction | 0.0151 | 0.2466 | 0.0127 – 0.0176 | 0.2070 – 0.2862 | **<0.001** |
| Aggregated real-life contacts | 0.1186 | 0.2037 | 0.0313 – 0.2060 | 0.0537 – 0.3537 | **0.008** |
| Momentary real-life contacts * Time point | -0.0073 | -0.0383 | -0.0189 – 0.0043 | -0.0993 – 0.0228 | 0.219 |
| Quality of most important interaction * Time point | 0.0005 | 0.0078 | -0.0018 – 0.0028 | -0.0295 – 0.0452 | 0.681 |
| **Random Effects** | | | | | |
| σ^2^ | 0.4215 | | | | |
| τ_00_ _Participants_ | 0.5022 | | | | |
| τ_11_ _Time of day_ | 0.0006 | | | | |
| τ_11_ _Real-life contacts_ | 0.0005 | | | | |
| τ_11_ _Quality of most important interaction_ | 0.0001 | | | | |
| ICC | 0.52 | | | | |
| N _VPNr_ | 70 | | | | |
| Observations | 5820 | | | | |
| Marginal R^2^ / Conditional R^2^ | 0.125 / 0.576 | | | | |

### Supplementary Table 3a.

Mixed model results for social contacts and negative affective states across both time points

|  | **Negative Affect** | | | | |
| --- | --- | --- | --- | --- | --- |
| *Predictors* | *Estimates* | *std. Beta* | *CI* | *standardized CI* | *p* |
| (Intercept) | 0.7908 | -0.1166 | 0.3841 – 1.1974 | -0.3139 – 0.0807 | <0.001 |
| Stressful life events | 0.0104 | 0.2886 | 0.0041 – 0.0166 | 0.1141 – 0.4630 | **0.001** |
| Psychosocial risk at birth | 0.0343 | 0.0785 | -0.0402 – 0.1088 | -0.0921 – 0.2491 | 0.367 |
| Gender | 0.1678 | 0.2138 | -0.0635 – 0.3990 | -0.0809 – 0.5086 | 0.155 |
| Time of day | -0.0028 | -0.0161 | -0.0071 – 0.0015 | -0.0409 – 0.0087 | 0.204 |
| Time point | 0.0348 | 0.0443 | 0.0102 – 0.0593 | 0.0131 – 0.0756 | **0.005** |
| Momentary real-life contacts | -0.0035 | -0.0214 | -0.0146 – 0.0076 | -0.0898 – 0.0471 | 0.540 |
| Aggregated real-life contacts | -0.0755 | -0.1636 | -0.1424 – -0.0086 | -0.3085 – -0.0187 | **0.027** |
| Momentary real-life contacts * Time point | 0.0023 | 0.0142 | -0.0060 – 0.0106 | -0.0369 – 0.0654 | 0.585 |
| **Random Effects** | | | | | |
| σ^2^ | 0.2384 | | | | |
| τ_00_ _Participants_ | 0.3073 | | | | |
| τ_11_ _Time of day_ | 0.0002 | | | | |
| τ_11_ _Real-life contacts_ | 0.0016 | | | | |
| ICC | 0.57 | | | | |
| N _VPNr_ | 70 | | | | |
| Observations | 6837 | | | | |
| Marginal R^2^ / Conditional R^2^ | 0.136 / 0.630 | | | | |

### Supplementary Table 3a.

Mixed model results for social contacts and negative affective states across both time points

|  | **Negative Affect** | | | | |
| --- | --- | --- | --- | --- | --- |
| *Predictors* | *Estimates* | *std. Beta* | *CI* | *standardized CI* | *p* |
| (Intercept) | 0.8142 | -0.1102 | 0.4488 – 1.1795 | -0.3040 – 0.0836 | <0.001 |
| Stressful life events | 0.0095 | 0.2847 | 0.0039 – 0.0151 | 0.1173 – 0.4520 | **0.001** |
| Psychosocial risk at birth | 0.0288 | 0.0698 | -0.0378 – 0.0953 | -0.0915 – 0.2311 | 0.396 |
| Gender | 0.1733 | 0.2349 | -0.0319 – 0.3785 | -0.0433 – 0.5132 | 0.098 |
| Time of day | -0.0019 | -0.0116 | -0.0059 – 0.0021 | -0.0357 – 0.0125 | 0.345 |
| Time point | 0.0179 | 0.0254 | -0.0064 – 0.0421 | -0.0080 – 0.0588 | 0.148 |
| Momentary real-life contacts | 0.0019 | 0.0130 | -0.0043 – 0.0081 | -0.0293 – 0.0554 | 0.547 |
| Quality of most important interaction | -0.0073 | -0.1545 | -0.0092 – -0.0053 | -0.1962 – -0.1128 | **<0.001** |
| Aggregated real-life contacts | -0.0657 | -0.1474 | -0.1246 – -0.0068 | -0.2796 – -0.0153 | **0.029** |
| Momentary real-life contacts * Time point | 0.0021 | 0.0142 | -0.0057 – 0.0098 | -0.0391 – 0.0676 | 0.601 |
| Quality of most important interaction * Time point | 0.0001 | 0.0013 | -0.0015 – 0.0016 | -0.0321 – 0.0347 | 0.938 |
| **Random Effects** | | | | | |
| σ^2^ | 0.1971 | | | | |
| τ_00_ _Participants_ | 0.2848 | | | | |
| τ_11_ _Time of day_ | 0.0002 | | | | |
| τ_11_ _Real-life contacts_ | 0.0003 | | | | |
| τ_11_ _Quality of most important interaction_ | 0.0001 | | | | |
| ICC | 0.57 | | | | |
| N _VPNr_ | 70 | | | | |
| Observations | 5820 | | | | |
| Marginal R^2^ / Conditional R^2^ | 0.155 / 0.640 | | | | |

### Supplementary Table 4a.

Mixed model results for social contacts, right amygdala volume, and positive affective state across both time points

|  | **Positive Affect** | | | | |
| --- | --- | --- | --- | --- | --- |
| *Predictors* | *Estimates* | *std. Beta* | *CI* | *standardized CI* | *p* |
| (Intercept) | 4.7536 | 0.0284 | 4.1717 – 5.3355 | -0.1893 – 0.2461 | **<0.001** |
| ICV | -0.0020 | -0.0438 | -0.0107 – 0.0066 | -0.2325 – 0.1449 | 0.649 |
| Stressful life events | 0.0015 | 0.1916 | -0.0001 – 0.0031 | -0.0126 – 0.3959 | 0.066 |
| Psychosocial risk at birth | -0.0412 | -0.0739 | -0.1420 – 0.0595 | -0.2545 – 0.1067 | 0.423 |
| Gender | 0.0410 | 0.0409 | -0.3470 – 0.4289 | -0.3463 – 0.4281 | 0.836 |
| Time of day | -0.0002 | -0.0011 | -0.0081 – 0.0076 | -0.0365 – 0.0344 | 0.952 |
| Time point | -0.0746 | -0.0734 | -0.1098 – -0.0394 | -0.1085 – -0.0383 | **<0.001** |
| Right amygdala volume | -0.0005 | -0.0967 | -0.0016 – 0.0006 | -0.3184 – 0.1250 | 0.393 |
| Momentary real-life contacts | 0.0214 | 0.1033 | 0.0086 – 0.0342 | 0.0417 – 0.1649 | **0.001** |
| Aggregated real-life contacts | 0.1389 | 0.2356 | 0.0478 – 0.2299 | 0.0811 – 0.3901 | **0.003** |
| Right Amygdala * Time point | -0.0002 | -0.0468 | -0.0004 – -0.0001 | -0.0814 – -0.0122 | **0.008** |
| Momentary real-life contacts * Time point | 0.0015 | 0.0075 | -0.0104 – 0.0134 | -0.0500 – 0.0649 | 0.807 |
| Momentary real-life contacts * right amygdala volume | -0.0000 | -0.0044 | -0.0001 – 0.0001 | -0.0668 – 0.0580 | 0.890 |
| Momentary real-life contacts * right amygdala volume * Time point | -0.0000 | -0.0139 | -0.0001 – 0.0001 | -0.0790 – 0.0511 | 0.675 |
| **Random Effects** | | | | | |
| σ^2^ | 0.4915 | | | | |
| τ_00_ _Participants_ | 0.5137 | | | | |
| τ_11_ _Time of day_ | 0.0009 | | | | |
| τ_11_ _Real-life contacts_ | 0.0017 | | | | |
| ICC | 0.51 | | | | |
| N _VPNr_ | 69 | | | | |
| Observations | 6837 | | | | |
| Marginal R^2^ / Conditional R^2^ | 0.092 / 0.546 | | | | |

### Supplementary Table 4b.

Mixed model results for social contacts, right amygdala volume, and positive affective state across both time points

|  | **Positive Affect** | | | | |
| --- | --- | --- | --- | --- | --- |
| *Predictors* | *Estimates* | *std. Beta* | *CI* | *standardized CI* | *p* |
| (Intercept) | 4.6419 | -0.0302 | 4.4153 – 4.8684 | -0.2445 – 0.1842 | <0.001 |
| ICV | 0.0014 | 0.1904 | -0.0001 – 0.0029 | -0.0139 – 0.3948 | 0.068 |
| Stressful life events | -0.0023 | -0.0505 | -0.0112 – 0.0065 | -0.2416 – 0.1406 | 0.605 |
| Psychosocial risk at birth | -0.0269 | -0.0500 | -0.1239 – 0.0701 | -0.2304 – 0.1303 | 0.587 |
| Gender | 0.1824 | 0.1887 | -0.1862 – 0.5509 | -0.1927 – 0.5701 | 0.332 |
| Time of day | -0.0030 | -0.0141 | -0.0103 – 0.0043 | -0.0479 – 0.0198 | 0.416 |
| Time point | -0.0988 | -0.1022 | -0.1348 – -0.0627 | -0.1400 – -0.0643 | **<0.001** |
| Right amygdala volume | -0.0006 | -0.1417 | -0.0017 – 0.0004 | -0.3658 – 0.0824 | 0.215 |
| Momentary real-life contacts | 0.0134 | 0.0701 | 0.0041 – 0.0228 | 0.0212 – 0.1189 | **0.005** |
| Quality of most important interaction | 0.0152 | 0.2469 | 0.0128 – 0.0176 | 0.2072 – 0.2866 | **<0.001** |
| Aggregated real-life contacts | 0.1213 | 0.2080 | 0.0341 – 0.2086 | 0.0584 – 0.3575 | **0.006** |
| Right Amygdala * Time point | -0.0003 | -0.0622 | -0.0004 – -0.0001 | -0.0996 – -0.0247 | **0.001** |
| Momentary real-life contacts * Time point | -0.0073 | -0.0371 | -0.0192 – 0.0046 | -0.0995 – 0.0252 | 0.231 |
| Momentary real-life contacts * right amygdala volume | -0.000003 | -0.0004 | -0.0000 – 0.0000 | -0.0527 – 0.0518 | 0.987 |
| Quality of most important interaction * Time point | 0.0005 | 0.0074 | -0.0018 – 0.0028 | -0.0304 – 0.0452 | 0.686 |
| Quality of most important interaction * right amygdala volume | 0.000002 | 0.0118 | -0.0000 – 0.0000 | -0.0297 – 0.0533 | 0.576 |
| Momentary real-life contacts * right amygdala volume * Time point | -0.00001 | -0.0165 | -0.0001 – 0.0001 | -0.0885 – 0.0555 | 0.654 |
| Quality of most important interaction * right amygdala volume * Time point | 0.000001 | 0.0080 | -0.0000 – 0.0000 | -0.0309 – 0.0468 | 0.688 |
| **Random Effects** | | | | | |
| σ^2^ | 0.4209 | | | | |
| τ_00_ _Participants_ | 0.5006 | | | | |
| τ_11_ _Time of day_ | 0.0007 | | | | |
| τ_11_ _Real-life contacts_ | 0.0005 | | | | |
| τ_11_ _Quality of most important interaction_ | 0.0001 | | | | |
| ICC | 0.51 | | | | |
| N _VPNr_ | 69 | | | | |
| Observations | 5742 | | | | |
| Marginal R^2^ / Conditional R^2^ | 0.143 / 0.581 | | | | |

### Supplementary Table 5.

Mixed model results for social contacts, left amygdala volume, and positive affective states across both time points

|  | **Positive Affect** | | | | |
| --- | --- | --- | --- | --- | --- |
| *Predictors* | *Estimates* | *std. Beta* | *CI* | *standardized CI* | *p* |
| (Intercept) | 4.7180 | -0.0297 | 4.1863 – 5.2497 | -0.2357 – 0.1764 | <0.001 |
| ICV | 0.0016 | 0.2189 | 0.0002 – 0.0031 | 0.0234 – 0.4143 | **0.028** |
| Stressful life events | -0.0012 | -0.0272 | -0.0092 – 0.0068 | -0.2101 – 0.1558 | 0.771 |
| Psychosocial risk at birth | -0.0245 | -0.0455 | -0.1181 – 0.0691 | -0.2192 – 0.1282 | 0.608 |
| Gender | 0.1730 | 0.1796 | -0.1647 – 0.5108 | -0.1709 – 0.5302 | 0.315 |
| Time of day | -0.0030 | -0.0141 | -0.0102 – 0.0041 | -0.0474 – 0.0193 | 0.409 |
| Time point | -0.0978 | -0.1021 | -0.1334 – -0.0623 | -0.1397 – -0.0646 | **<0.001** |
| Left amygdala volume | -0.0010 | -0.1950 | -0.0021 – -0.0000 | -0.3927 – 0.0027 | **0.049** |
| Momentary real-life contacts | 0.0138 | 0.0713 | 0.0052 – 0.0223 | 0.0265 – 0.1160 | **0.002** |
| Quality of most important interaction | 0.0152 | 0.2472 | 0.0128 – 0.0176 | 0.2075 – 0.2869 | **<0.001** |
| Aggregated real-life contacts | 0.0994 | 0.1708 | 0.0141 – 0.1848 | 0.0243 – 0.3173 | **0.022** |
| Left Amygdala * Time point | -0.0003 | -0.0737 | -0.0005 – -0.0002 | -0.1110 – -0.0364 | **<0.001** |
| Momentary real-life contacts * Time point | -0.0066 | -0.0321 | -0.0182 – 0.0049 | -0.0928 – 0.0286 | 0.260 |
| Momentary real-life contacts * Left amygdala volume | 0.00003 | 0.0358 | -0.0000 – 0.0001 | -0.0120 – 0.0836 | 0.142 |
| Quality of most important interaction * Time point | 0.0004 | 0.0070 | -0.0019 – 0.0027 | -0.0304 – 0.0444 | 0.717 |
| Quality of most important interaction * Left amygdala volume | 0.000001 | 0.0150 | -0.0000 – 0.0000 | -0.0261 – 0.0562 | 0.474 |
| Momentary real-life contacts * Left amygdala volume * Time point | -0.0001 | -0.0900 | -0.0002 – -0.0000 | -0.1552 – -0.0247 | **0.007** |
| Quality of most important interaction * Left amygdala volume * Time point | -0.000001 | -0.0040 | -0.0000 – 0.0000 | -0.0430 – 0.0349 | 0.840 |
| **Random Effects** | | | | | |
| σ^2^ | 0.4201 | | | | |
| τ_00_ _Participants_ | 0.4843 | | | | |
| τ_11_ _Time of day_ | 0.0007 | | | | |
| τ_11_ _Real-life contacts_ | 0.0004 | | | | |
| τ_11_ _Quality of most important interaction_ | 0.0001 | | | | |
| ICC | 0.50 | | | | |
| N _VPNr_ | 70 | | | | |
| Observations | 5820 | | | | |
| Marginal R^2^ / Conditional R^2^ | 0.159 / 0.582 | | | | |

### Supplementary Table 6.

Mixed model results for social contacts, neuroticism, and positive affective states across both time points

|  | **Positive Affect** | | | | |
| --- | --- | --- | --- | --- | --- |
| *Predictors* | *Estimates* | *std. Beta* | *CI* | *standardized CI* | *p* |
| (Intercept) | 4.3638 | -0.0178 | 3.8327 – 4.8950 | -0.2041 – 0.1686 | <0.001 |
| Stressful life events | 0.0048 | 0.1109 | -0.0037 – 0.0134 | -0.0861 – 0.3080 | 0.270 |
| Psychosocial risk at birth | -0.0561 | -0.1067 | -0.1449 – 0.0327 | -0.2754 – 0.0621 | 0.216 |
| Gender | 0.1368 | 0.1449 | -0.1365 – 0.4101 | -0.1446 – 0.4345 | 0.327 |
| Time of day | -0.0029 | -0.0136 | -0.0102 – 0.0044 | -0.0482 – 0.0210 | 0.442 |
| Time point | -0.0865 | -0.0970 | -0.1220 – -0.0510 | -0.1353 – -0.0587 | **<0.001** |
| Neuroticism | -0.0337 | -0.3280 | -0.0507 – -0.0166 | -0.4933 – -0.1627 | **<0.001** |
| Momentary real-life contacts | 0.0161 | 0.0831 | 0.0063 – 0.0259 | 0.0328 – 0.1334 | **0.001** |
| Quality of most important interaction | 0.0151 | 0.2519 | 0.0126 – 0.0176 | 0.2106 – 0.2932 | **<0.001** |
| Aggregated real-life contacts | 0.0780 | 0.1327 | -0.0050 – 0.1610 | -0.0085 – 0.2739 | 0.066 |
| Neuroticism * Time point | 0.0051 | 0.0407 | 0.0012 – 0.0090 | 0.0029 – 0.0786 | **0.011** |
| Momentary real-life contacts * Time point | -0.0120 | -0.0606 | -0.0240 – 0.0001 | -0.1225 – 0.0013 | 0.051 |
| Momentary real-life contacts * Neuroticism | -0.0002 | -0.0117 | -0.0014 – 0.0009 | -0.0653 – 0.0419 | 0.668 |
| Quality of most important interaction * Time point | 0.0006 | 0.0097 | -0.0017 – 0.0029 | -0.0286 – 0.0479 | 0.630 |
| Quality of most important interaction * Neuroticism | 0.0001 | 0.0183 | -0.0001 – 0.0004 | -0.0217 – 0.0583 | 0.369 |
| Momentary real-life contacts * Neuroticism * Time point | -0.0022 | -0.1062 | -0.0037 – -0.0008 | -0.1732 – -0.0391 | **0.002** |
| Quality of most important interaction * Neuroticism * Time point | -0.0002 | -0.0339 | -0.0004 – 0.0000 | -0.0688 – 0.0009 | 0.056 |
| **Random Effects** | | | | | |
| σ^2^ | 0.4089 | | | | |
| τ_00_ _Participants_ | 0.3882 | | | | |
| τ_11_ _Time of day_ | 0.0007 | | | | |
| τ_11_ _Real-life contacts_ | 0.0007 | | | | |
| τ_11_ _Quality of most important interaction_ | 0.0001 | | | | |
| ICC | 0.47 | | | | |
| N _VPNr_ | 69 | | | | |
| Observations | 5742 | | | | |
| Marginal R^2^ / Conditional R^2^ | 0.183 / 0.569 | | | | |

### Supplementary Table 7.

Mixed model results for social contacts, SCZ-PRS, and positive affective states across both time points

|  | **Positive Affect** | | | | |
| --- | --- | --- | --- | --- | --- |
| *Predictors* | *Estimates* | *std. Beta* | *CI* | *standardized CI* | *p* |
| (Intercept) | 4.5638 | -0.0589 | 3.9908 – 5.1368 | -0.2697 – 0.1519 | <0.001 |
| Stressful life events | 0.0002 | 0.0046 | -0.0084 – 0.0088 | -0.1929 – 0.2022 | 0.964 |
| Psychosocial risk at birth | -0.0833 | -0.1551 | -0.1929 – 0.0262 | -0.3589 – 0.0487 | 0.136 |
| Gender | 0.2451 | 0.2533 | -0.0797 – 0.5699 | -0.0824 – 0.5890 | 0.139 |
| Time of day | -0.0020 | -0.0094 | -0.0093 – 0.0052 | -0.0428 – 0.0241 | 0.584 |
| Time point | -0.1047 | -0.1081 | -0.1408 – -0.0686 | -0.1462 – -0.0700 | **<0.001** |
| SCZ-PRS | -0.0765 | -0.0836 | -0.2389 – 0.0859 | -0.2559 – 0.0887 | 0.356 |
| Momentary real-life contacts | 0.0135 | 0.0719 | 0.0043 – 0.0226 | 0.0234 – 0.1204 | **0.004** |
| Quality of most important interaction | 0.0151 | 0.2481 | 0.0127 – 0.0175 | 0.2080 – 0.2883 | **<0.001** |
| Aggregated real-life contacts | 0.1022 | 0.1753 | 0.0007 – 0.2036 | 0.0012 – 0.3494 | **0.048** |
| SCZ-PRS * Time point | -0.0732 | -0.0817 | -0.1075 – -0.0389 | -0.1187 – -0.0447 | **<0.001** |
| Momentary real-life contacts * Time point | -0.0027 | -0.0135 | -0.0149 – 0.0095 | -0.0782 – 0.0511 | 0.663 |
| Momentary real-life contacts * SCZ-PRS | -0.0053 | -0.0289 | -0.0137 – 0.0031 | -0.0744 – 0.0167 | 0.214 |
| Quality of most important interaction * Time point | 0.0004 | 0.0068 | -0.0019 – 0.0027 | -0.0309 – 0.0446 | 0.741 |
| Quality of most important interaction * SCZ-PRS | 0.0013 | 0.0223 | -0.0011 – 0.0037 | -0.0178 – 0.0623 | 0.275 |
| Momentary real-life contacts * SCZ-PRS * Time point | -0.0088 | -0.0481 | -0.0188 – 0.0012 | -0.1025 – 0.0063 | 0.083 |
| Quality of most important interaction * SCZ-PRS * Time point | -0.0018 | -0.0309 | -0.0039 – 0.0003 | -0.0661 – 0.0043 | 0.085 |
| **Random Effects** | | | | | |
| σ^2^ | 0.4179 | | | | |
| τ_00_ _Participants_ | 0.4835 | | | | |
| τ_11_ _Time of day_ | 0.0006 | | | | |
| τ_11_ _Real-life contacts_ | 0.0005 | | | | |
| τ_11_ _Quality of most important interaction_ | 0.0001 | | | | |
| ICC | 0.51 | | | | |
| N _VPNr_ | 68 | | | | |
| Observations | 5631 | | | | |
| Marginal R^2^ / Conditional R^2^ | 0.188 / 0.604 | | | | |
